# Individual Differences in Audio-Visual Binding Can Predict the Varied Severity of Motion Sickness

**DOI:** 10.1101/2022.05.09.491170

**Authors:** Revital Zilka, Yoram Bonneh

## Abstract

In neuroscience, research often focuses on the group, while ignoring large individual differences, which are left poorly understood. One such example is the large individual variability in the susceptibility to motion-sickness (MS), the feeling of sickness that typically occurs during travel or movement. Current explanations of MS focus on the sensory conflict in the perception of motion, primarily between the vestibular and the visual systems, e.g., when feeling motion but not seeing it. To account for the large individual differences in MS, we hypothesized that people feel motion sickness only when the conflicting stimuli are perceived as bound together, and their tendency to bind conflicting multi-sensory information is an individual trait. To test this hypothesis, we measured the persistence of audio-visual binding using the McGurk effect in which watching a moving mouth alters the auditory perception of phonemes. We used a temporal audio-visual mismatch to probe the persistence of binding and computed a temporal binding window for each individual (n=21) in 3 tasks: syllable Identification (McGurk), simultaneity judgement, and syllable synchronization judgement. To assess the severity of MS, we used 2 subjective symptom questionnaires. We found that the temporal binding window of the McGurk stimuli in two of the tasks varied across individuals and was positively correlated (R>0.8) with the MS questionnaire severity scores. These results support our hypothesis and shed new light on the enigmatic differences between individuals regarding their susceptibility to motion-sickness. They also highlight the potential strength of studies focusing on individual differences and neurological diversity.

**Significance Statement:** Motion sickness is the feeling of sickness that typically occurs during travel or movement. Although it is commonly linked to a sensory mismatch, the large differences in its susceptibility across people remains poorly understood. In the current study, we link these individual differences to a multi-modal temporal binding window, which measures the persistence of perceiving multi-modal stimuli as coming from the same source despite temporal discrepancies. We demonstrated this link by finding a high correlation between the temporal binding window of an audio-visual illusion (the McGurk effect) and the subjective reports of the susceptibility to motion sickness. These results shed new light on the enigmatic differences between individuals regarding their susceptibility to motion sickness and highlight the potential strength of studies focusing on individual differences and neurological diversity.

## Introduction

Motion sickness (MS) refers to the feeling of sickness that typically occurs during travel or movement. Nausea and vomiting are the primary symptoms of motion sickness; however, there is also a general weakness, general discomfort, stomach awareness, cold sweating, dizziness, headache, pallor, a sensation of bodily warmth, and more (J.F. Golding, 2016). Many theories and explanations have been suggested for MS; most of them are related to vestibular functions and the idea of a sensory conflict (J.F. Golding, 2016), commonly assumed to exist between the visual and vestibular signals of motion (Benson, 2002). Individuals vary widely in their susceptibility to MS. Currently there is evidence that genetic factors (Klosterhalfen et al., 2005; Stern, Hu, LeBlanc, & Koch, 1993), age (Reason, J. T., & Brand, 1975), and gender (RS Kennedy, DS Lanham, CJ Massey, 1995) affect the susceptibility to MS.

Multisensory integration is the process of combining sensory information obtained from multiple sensory modalities into a coherent percept. It enables appropriate behavioral responses to be created in circumstances where information from a single sensory modality is insufficient; therefore, it is important for survival (Sarko, Ghose, & Wallace, 2013). A key process in integrating multisensory information is cross-modal binding, which refers to the binding of information coming from different modalities such as audio and vision into one perceptual source (Murray, Lewkowicz, Amedi, & Wallace, 2016). One of the most striking cross-modal illusions is the McGurk effect, which is an audio-visual integration illusion (Mcgurk & Macdonald, 1976). It occurs when one hears an auditory syllable together with an incongruent visual syllable, yielding the perception of a third syllable (Mcgurk & Macdonald, 1976).

Since stimuli that come from different sensory modalities and at slightly different times can nevertheless be perceived as simultaneous or bound together and associated with the same event, one could investigate the temporal window in which this binding occurs; this is termed the “Temporal Binding Window” (TBW). This concept was extensively studied by Wallace and colleagues (Feldman et al., 2018; Powers, Hillock, & Wallace, 2009; Stevenson, Ryan A and Wallace, 2013; Stevenson, Ghose, et al., 2014; Stevenson, Zemtsov, & Wallace, 2012; van Wassenhove, Grant, & Poeppel, 2007; Wallace & Stevenson, 2014) The TBW could also be measured for the McGurk effect (van Wassenhove et al., 2007) by investigating the effect of Audio-Visual asynchrony on the identification of incongruent (McGurk) Audio-Visual speech stimuli, i.e., the TBW in which the illusion persists (van Wassenhove et al., 2007).

The decision as to whether to bind or not to bind multisensory information depends on deciding whether the information comes from the same source or perceptual object (Stevenson et al., 2012). For that purpose, the brain needs to examine the input in three dimensions: Spatial, Temporal, and Contextual congruency, assuming that sensory input from different sensory modalities that come from the same location in space, at the same time, and in contextual congruency come from the same source (Burr & Alais, 2006; Welch & Warren, 1980).

Could Motion Sickness be related to the TBW? The current study is based on a novel hypothesis that MS is related to the strength of the multisensory binding that can be measured by the TBW. This hypothesis is partly motivated by the similarity between the effect of age on the audio-visual TBW and motion sickness: both show a similar U-shaped function (J.F. Golding, 2016). To test this hypothesis, we conducted a set of experiments that measured the audio-visual TBW with the McGurk effect and correlated these measures across individuals with the strength of MS assessed by questionnaires.

## Materials and Methods

### Participants

Twenty-one healthy adults with normal or corrected to normal vision and without known hearing problems participated in the experiments.

### Stimuli and Procedures

The subjects first completed two questionnaires that estimate their susceptibility to MS, and then participated in four experiments that measured the Temporal Binding Window (TBW) via the effect of cross-modal asynchrony on McGurk stimuli perception. Since previous research suggests that the TBW depends on the exact task applied (Basu Mallick, F. Magnotti, & S. Beauchamp, 2015; Freeman et al., 2013), we tested three different tasks, all involving the McGurk stimuli: The classical McGurk effect with syllable identification (6 syllables), simultaneity judgement (Stevenson, Ryan A and Wallace, 2013; van Wassenhove et al., 2007), and syllable synchronization judgement. For the first two tasks, the stimuli were presented using a VLC media player on a 14” HD LCD monitor, with the visual stimulus occupying a window of around 15×15 deg, whereas the last task was implemented using Psych Toolbox.3 and code written in MATLAB.

### The McGurk Syllable Identification Task (McG)

To test the effect of temporal asynchrony on the McGurk effect, we used 43 short video clips (4 seconds per video); each video was a combination of an audio track that plays the BA syllable and a visual track that illustrates the pronunciation of the GA syllable. The relative time of the visual and auditory tracks was systematically manipulated across the clips, from perfect simultaneity to large asynchrony at 1000ms, with 50ms steps; in 20 videos the audio track preceded the visual (-SOA), and in 20 videos the visual preceded the audio track (+SOA). The order of trial presentation was from 0ms to 1000ms and then from −50ms to −1000ms. There were three additional conditions of a zero-lag video, an audio only video and a visual only video. The clips were created from a popular McGurk demo clip. Each clip had the same syllable repeated 6 times appearing via spacing as 3 ‘BA-BA’ pairs, with a total duration of 4 sec (McG). The participant was instructed to report for each video all 6 perceived syllables in order, selected from the 3 options of BA GA or DA. These reports were recorded.

### The Simultaneity Judgement Task (Time Sync)

For the simultaneity judgment task, we created a shorter 2-syllable stimulus by extracting the first 2 syllables of the McG video clip. We added 6 more SOAs to increase the accuracy: +/- 10ms, +/-20ms, +/-80ms, and excluded the audio only and visual only videos. The participant’s task was to report whether the auditory and visual stimuli were simultaneous in each trial. Note that this task does not inquire about speech perception—only about the timing of the auditory and visual stimuli.

### The Syllable Synchronization Judgement Task (Syllable Sync)

In this task, the participant was instructed to report for each video whether the syllable that they heard is phonetically the same syllable that they saw. The videos were presented in a pseudorandom order.

### Questionnaires for Assessing Motion Sickness

Each participant completed two questionnaires designed to quantify his/her susceptibility to MS. The first questionnaire was developed by us, to fit Israeli society, by emphasizing bus and private car travel but not boats and aircraft. The scores of this questionnaire (“MSDQ = Motion Sickness Driving Questionnaire”) ranged from 20 to 100 points; the threshold for MS was set to 30. For correlations, we converted the MSDQ to log units, since this appears to handle properly its repetitive structure. The second questionnaire is called the Motion Sickness Susceptibility Questionnaire Short-form (MSSQ-Short). The scores in this questionnaire ranged from 0 to 54 points, with a threshold for MS in adults set to 5.11 (John F. Golding, 1998). The participants first completed the questionnaires and then performed the McGurk binding experiment.

### Data Analysis

For each subject and task, the TBW was estimated in two ways. In the main way, the proportions of perceived “DA” or “in sync” (depending on the experiment) were computed as a function of SOA, then “smoothed” by convolving with a uniform kernel of 10 samples, followed by finding the 2 crossing points of the 0.5 (50%) threshold via interpolation, and subtracting the points to obtain the TBW in milliseconds (Figure 1). In a second method, a crude estimate, which is independent of smoothing, was extracted as the accumulated number of trials with reported “DA” or “in sync” per participant and task. In both methods, participants who did not perceive the McGurk effect under any condition were assigned a zero TBW.

**Fig. 1.**
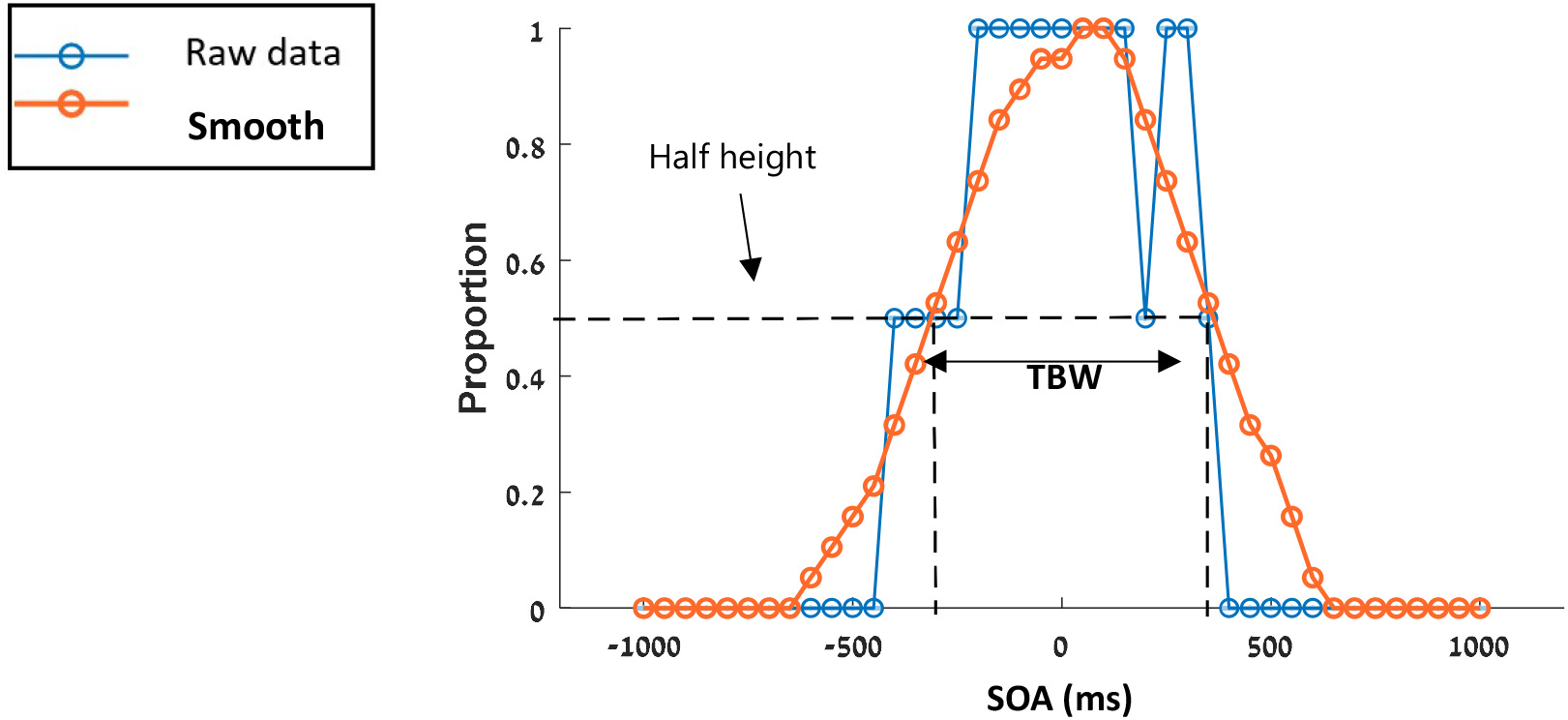
The method for computing the TBW in the McGurk experiments, exemplified on data from one participant. As shown, the raw data (blue) are not smooth due to the few trials collected per SOA (1 or 2 trials), making it necessary to smooth the data (see the text for more details). Then we find the 2 crossing points of the 0.5 (50%) threshold via interpolation and subtract the points to obtain the TBW in milliseconds. An alternative method that does not depend on smoothing, namely, summing all the data points, was also investigated.

## Results

We assessed the audio-visual TBW in 3 experiments: syllable identification, simultaneity judgement, and syllable synchronization judgement. The group averages for the perceptual reports in the McGurk syllable identification experiment with 6 syllables (McG) are shown in Figure 2(A). As shown, the frequency of perceiving DA peaks at an SOA of around +100ms and decreases gradually with asynchrony, reflecting the width of the temporal binding window. The perception of BA shows the opposite effect (a minimum at an SOA of ∼+100), and there is a small effect of GA with a time course similar to DA. These results are similar to those obtained in a previous study (van Wassenhove et al., 2007). The group averages for all perceptual reports in the McGurk binding experiment (McG, Syllable sync, and Time sync) are shown in Figure 2(B). As shown, the frequency of perceiving DA or sync in the syllable and time experiments peaks at an SOA of ∼+100 and decreases gradually with asynchrony, reflecting the width of the temporal binding window.

**Fig. 2.**
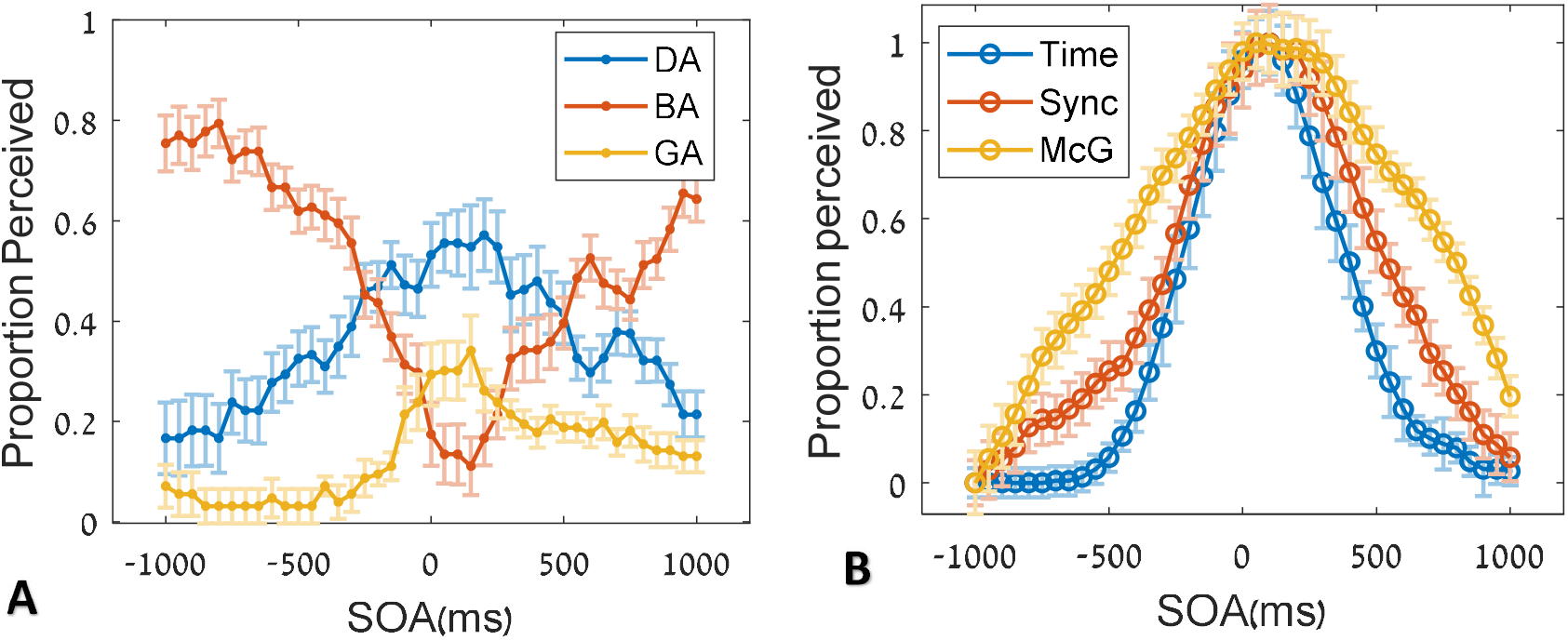
Group averages of the perceptual judgments as a function of audio-visual asynchrony (SOA). (*A*) The proportion of perceiving DA, BA, and GA. (*B*) Comparison of different tasks: the proportion of perceiving DA (McG), “the same syllable” (Syllable), or simultaneous presentation (Time), normalized by “clamping” the peaks to 1 to allow a comparison of tuning width. Data were averaged across observers, with error bars showing normalized standard error (SE). Note the difference in the binding width between the experiments.

Figure 3 shows six representative individual examples of audio-visual binding curves, 3 with narrow TBW: 504-728ms in McG, 537-658ms in Syllable Sync, and 488-561ms in Time Sync, as well as three examples with wide TBWs: 1122-1722ms in McG, 1366-1658ms in Syllable Sync, and 561-927ms in Time Sync. The individuals with narrow TBW also reported low MS in the questionnaires (MSSQ John F. Golding, 2006) and those with a wide TBW reported a high MS (John F. Golding, 2006).

**Fig. 3.**
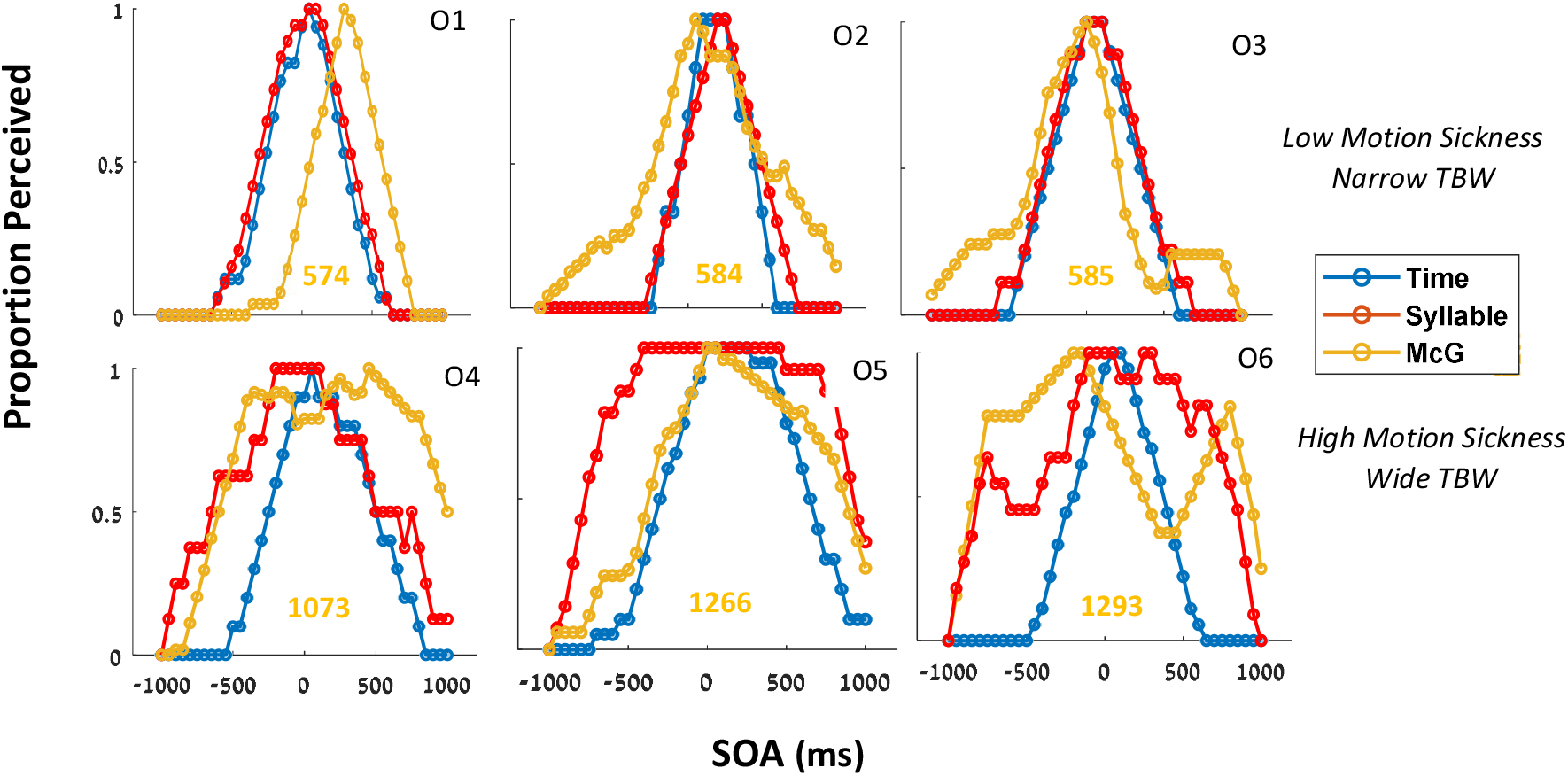
Individual audio-visual binding curve examples for the 3 tasks. The proportion of perceiving audio-visual binding is plotted as a function of SOA, with data smoothed (see the Methods). Top row: examples of individuals with a narrow temporal binding window (TBW) in the McGurk experiment and without motion sickness (by self-reports). Bottom row: examples of individuals with a wide TBW, and with motion sickness. The three tasks are color-coded as in Figure 2. The TBW was computed at a 50% cutoff following smoothing, with the TBW values in milisec for the “McG identification task”, shown in orange.

Figure 4 presents the correlation matrix of all TBWs in the three main experiments and the two MS questionnaires. The significant correlations are denoted in red, whereas non-significant correlations are in black. The correlation R is shown, and the level of significance is denoted by stars (see Figure 4, caption). In the correlations, there were N=21 subjects, except for the syllable sync and time sync experiments, both with N=16. The reduced N was partly due to the exclusion of cases with zero TBW, i.e., assigned when the McGurk effect was not perceived under any conditions.

**Fig. 4.**
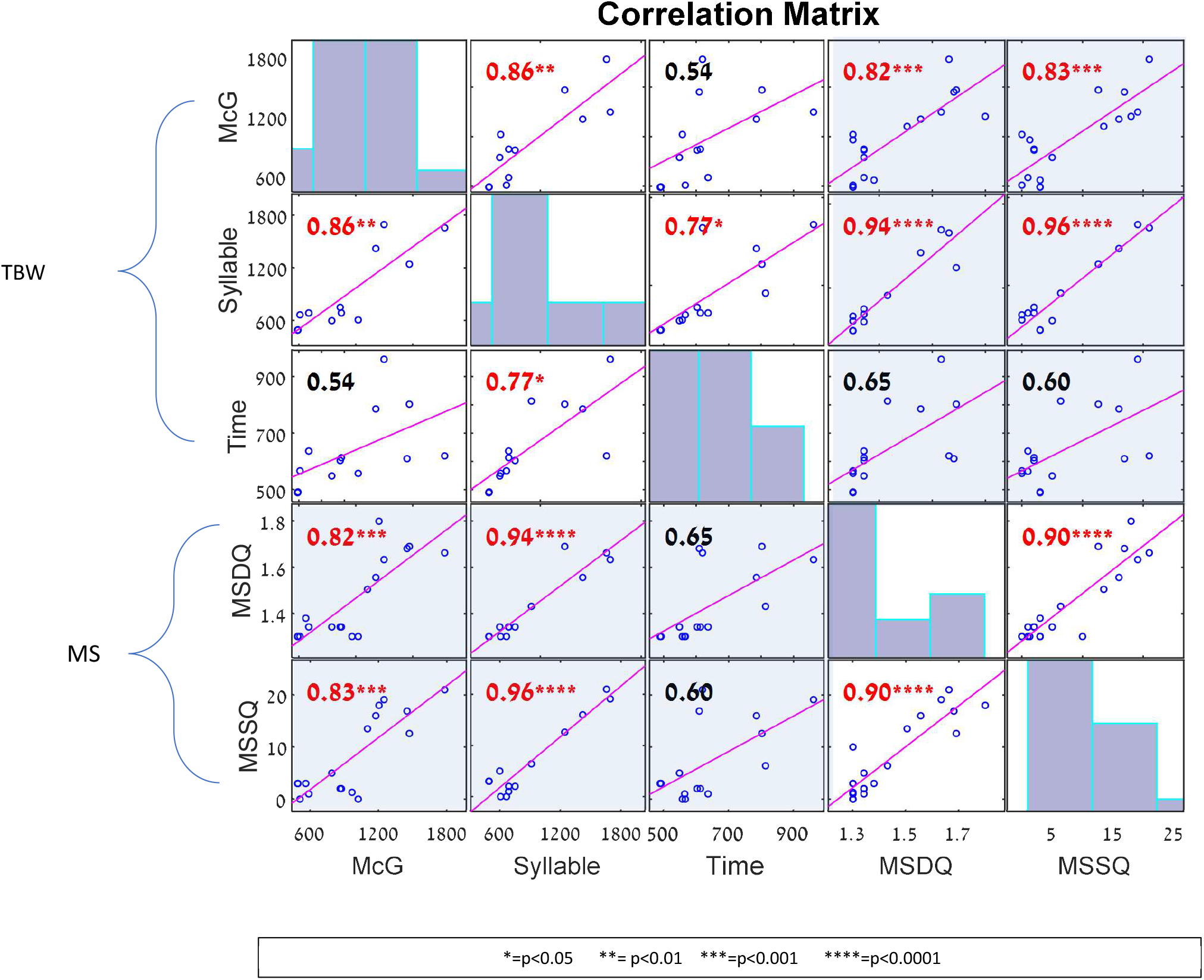
A correlation matrix of all correlations (R) between all TBWs in the three main experiments and the 2 questionnaires. The correlations related to the connection between Motion Sickness and audio-visual integration are denoted in light blue. The blue bar graphs on the diagonal plot the distribution histogram of the data on the X-axis. The number of participants was McGurk N=21, MSDQ N=21, Time sync and syllable sync N= 16, and MSSQ questionnaire N= 21. The highest correlations were found between Syllable sync and MSSQ: R=0.96, between Syllable sync and MSDQ: R=0.94, between McG and MSDQ: R=0.82 and between McG and MSSQ: R=0.83. The statistical significance marks (stars) were determined by p-values corrected for multiple comparisons (Bonferoni) assuming 10 comparisons.

All the significant p-values were corrected for multiple comparisons, via a conservative estimate (Bonferroni) and assuming that there were 10 comparisons (Figure 4). Some of the correlations examine the consistency of the measures and they appear in a white background. The two MS questionnaires were highly correlated (R=0.9, p<0.0001). Among the 3 TBW measures, the McG and Syllable sync were significantly correlated (R=0.86, p<0.01). The correlation between the TBW measures and the questionnaires (Figure 4, shaded plots) was found to be significant for the McG and Syllable sync, but not for the Time sync. The highest correlation was found between the Syllable sync and the MSSQ questionnaire (R=0.96, p<0.0001). Table 1 presents all the correlation values and their uncorrected p-values from Figure 4. We also examined an alternative estimate of the audio-visual integration based on a summation of the raw data without explicitly calculating the TBW. The results were similar to the calculated TBW, although the correlations were lower. For example, the correlation between MSSQ and the McGurk (a summation of raw data) was 0.7.

**Table 1.**
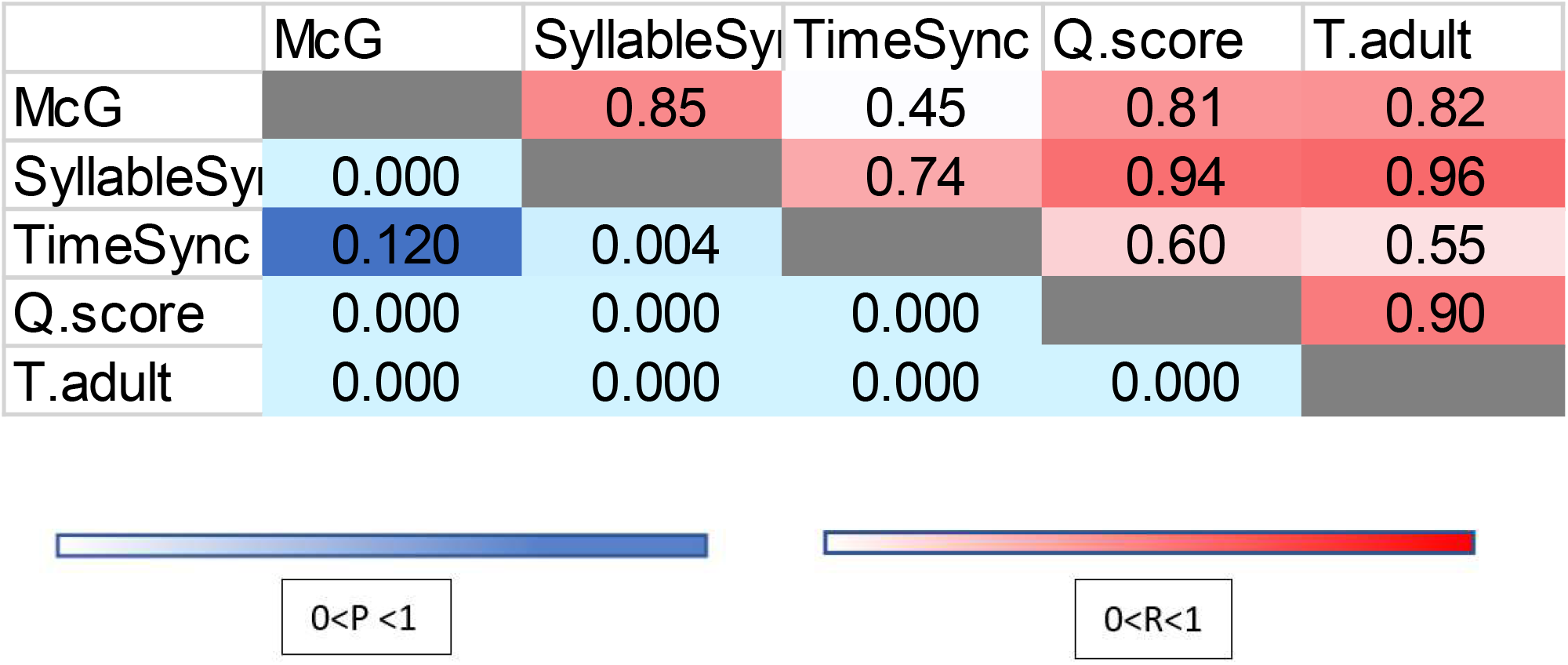
A correlations table that shows the correlations (R) between all TBWs in the three major experiments and the 2 questionnair

When examining the MS questionnaire scores, one can notice a cluster of people with very little MS and others who have higher degree of it (Figure 5A). We therefore divided the subjects into two groups via a questionnaire threshold and analyzed the difference in TBW between groups. The results are shown in Figure 5; the clustering for the two questionnaires is shown in Figure 5A, divided into the two clusters via a threshold. This threshold was derived from a previous work for the MSSQ (John F. Golding, 1998), and manually selected to divide the sample equally for the MSDQ. Using the MSSQ threshold, we analyzed the TBW for syllable sync (Figure 5B) and the TBW for McGurk (Figure 5C) with a bee swarm and distribution plots. As shown, the difference in TBWs between groups was highly significant, with both showing no overlap, i.e., AUC=1 (the area under the ROC curve), and a large effect size (2.8 and 3.6 for the Syllable and McG TBWs, respectively).

**Fig. 5.**
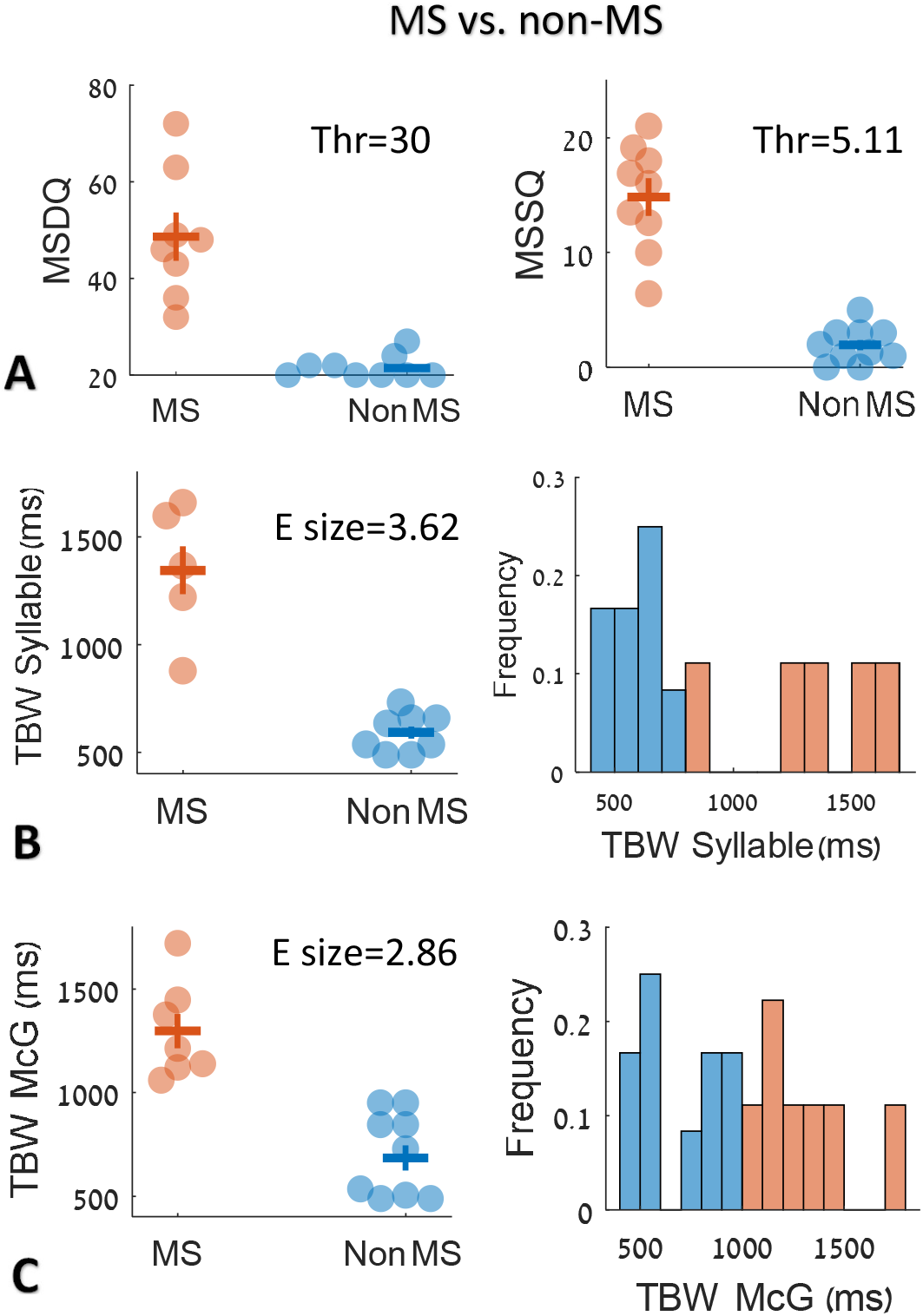
Motion sickness results by groups. Using the questionnaires, the subjects were divided into high and low MS by a threshold, and the results were compared in a bee swarm plot. (A) The questionnaire scores, MSDQ (our adapted questionnaire), and MSSQ for adults (Golding, 2006). (B) TBW for the syllable task, and (C) TBW for the McG task. As shown, dividing the participants into low (n=9) and high (n=7) MS according to their self-reports in the MSSQ questionnaire resulted in a perfect dissociation (AUC=1) for the two TBW measures. The average TBW-syllable was ∼1350ms and ∼650ms for high and low MS, respectively, with an effect size of 3.62.

## Discussion

In the present study, we attempted to better understand the individual differences in the severity of motion sickness (MS). Our research question focused on the connection between audio-visual binding and the severity of MS. We hypothesized that the width of the binding window in audio-visual integration is correlated with the susceptibility to MS, with narrow binding associated with low susceptibility and wide binding associated with high susceptibility. The idea behind this hypothesis is that there are common properties of multisensory integration across modalities. Accordingly, the tendency to bind or fuse rather than dissociate audiovisual stimuli will be similar to the tendency to bind or fuse vestibular-visual stimuli. In addition, we assumed that MS results from a conflict in the multisensory integration between the vestibular and visual stimuli (J.F. Golding, 2016) and that this conflict is eliminated or ignored when binding is prevented. Our results confirm this hypothesis by showing a correlation between the audio-visual TBW and MS severity assessed by questionnaires. Next, we will discuss the robustness of the findings and their interpretation, as well as additional preliminary experiments with physiological measures.

### The Robustness of the Findings

In designing our study, we took several measures to ensure the robustness of our findings. Basu Mallick et al. (2015) reported a high variability in people’s response to different videos of the McGurk effect, due to a difference in the person speaking or the number of repetitions in the video. Therefore, in our study we repeated the experiments using different speakers as well as a different number (2,6) of syllable repetitions (Basu Mallick et al., 2015).

In our results (Figure 2) we found asymmetries between audio lag and audio lead, as previously reported (Stevenson, Ryan A and Wallace, 2013). Wallace and his colleagues found that people are more likely to perceive stimuli synchronously when the visual stimulus precedes the auditory stimulus in time. One explanation for this effect is that in a natural environment the visual stimulus reaches the retina before the auditory stimulus reaches the cochlea (E.Poppel, 1990). To account for individual differences in MS, we averaged the audio lag and lead measures to obtain more statistical power and accuracy and left the analysis of this asymmetry and its possible relation to MS for future work.

Another important measure we took to achieve robustness as well as a deeper understanding of the individual differences in audio-visual integration was to use three different tasks. Previous studies compared syllable identification (McGurk) to subjective simultaneity judgment (time synchronization) (Stevenson, Ryan A and Wallace, 2013; van Wassenhove et al., 2007), but we are the first to apply a third task of “phonetic synchronization”. We suspected that a person might be able to notice a delay in time between audio-visual stimuli, but at the same time, fail to notice the phonetic discrepancy, and indeed this was the case in our experiments.

### The effect of age and gender

An interesting support of our hypothesis can be derived from the studies on TBW across ages. Whereas during development the TBW narrows (Sarko, Nidiffer, et al., 2013), this effect reverses in older (>60 years) people (Chan, Pianta, & McKendrick, 2014). This U-shaped effect appears to be similar to the susceptibility to MS, with very young and old people more susceptible (J.F. Golding, 2016; Turner & Griffin, 1999). Reason and Brand (Reason, J. T., & Brand, 1975) claimed that infants and toddlers up to 2 years old are immune to MS, even though they have no vomiting immunity. They also believe that MS begins around age 6-7, but sometimes even before, and that the peak susceptibility is around 9 years old (Henriques, Douglas de Oliveira, Oliveira-Ferreira, & Andrade, 2014; Turner & Griffin, 1999). Then there is another decrease in sensitivity during adulthood up to the age of 20, and evidence, though limited, for a gradual decrease in sensitivity until old age; a minority of cases overcame MS towards old age (J.F. Golding, 2016). In our study we examined participants around the age of 30 to 40.

There is also evidence of a gender effect in motion sickness. Sea, dry, and air transportation surveys indicate that women are more susceptible to traffic sickness and have a higher incidence of vomiting and nausea (RS Kennedy, DS Lanham, CJ Massey, 1995). In the current study we had 5 females (out of 12) with MS and only 2 males (out of 8) with MS, i.e., there was a 5:3 ratio between females and males with MS.

### Comparison Between the Different TBW Measures

In comparing the different TBW measures, we found that the TIME TBW was the narrowest, and that the McG TBW was the widest, with SYLLABLE TBW in the middle. We found that most participants were able to detect more easily time delays than phonetic synchronization (i.e., consistency). That is, they were able to easily notice that there were time delays between the auditory and visual stimuli, but at the same time, continued to perceive them as phonetically synchronized. The difference between these two measures was significant and greater among participants with motion sickness, who have a difference of about 676ms between the phonetic and temporal synchronization windows, compared with a difference of about 55ms in non-motion sickness participants. In general, the dependency of the TBW on the task in our results is consistent with previous studies of multisensory integration (Freeman, 2016; Freeman et al., 2013; Ipser et al., 2017; Ipser, Karlinski, & Freeman, 2018),

### Motion Sickness and Audio-Visual Binding

The main finding of the current study is the high correlation between the Motion Sickness questionnaires (MSDQ, our MS questionnaire, or the MSSQ adult (John F. Golding, 2006)) and the McG TBW (R = 0.81 and 0.82, respectively) as well as the Syllable Sync TBW (R=0.94 and 0.96, respectively, see Figure 4). Based on these results, we suggest that people differ in the way they perform multisensory integration, and that this relates to MS in one of two ways: (1) Strong (or wide) binding is associated with high susceptibility to MS; (2) Weak (or narrow) binding is associated with low susceptibility to MS. We hypothesize that this property should generalize to all sensory modalities.

The results of the current study could shed light on the susceptibility to MS in clinical populations such as Autism Spectrum Disorder (ASD). Stevenson et al (Stevenson, Siemann, et al., 2014) found a wider audio-visual TBW for speech sounds in children and teenagers with high functioning ASD (but their TBW measure could be different from our McGurk-based TBW). According to the current findings, a wider TBW predicts a higher susceptibility to MS in ASD. Currently, we could not find any quantitative assessments of MS in ASD. We are aware of some anecdotal evidence for increased MS in ASD, as well as some evidence for the opposite, i.e., lack of MS in ASD. Noteworthy, Ritvo and Ornitz (Ornitz, 1970; Ritvo et al., 1969) found **d**ecreased post-rotatory nystagmus in early infantile autism, indicative of less MS in conditions that typically induce it. We are aware of a tendency in the non-verbal severely autistic to dissociate between the senses or shutdown one sense (see (Bonneh et al., 2008) for a case study) which predicts less MS in severe autism. More research is needed to clear this issue.

## Conclusions

Motion sickness varies across people and is typically explained in terms of a physiological response to a perceptual conflict between the visual and vestibular modalities. The tendency to ignore a multi-sensory mismatch, presumably by “shutting off” one sensory input, predicts the severity of motion sickness, as we found in the current study for audio-visual binding. This tendency is likely to generalize across modalities and may reflect a currently unknown “perceptual style” or even specific personality traits.

